# An energy coupling factor transporter of *Streptococcus sanguinis* impacts antibiotic susceptibility as well as metal and membrane homeostasis

**DOI:** 10.1101/2024.07.12.603315

**Authors:** Marta Rudzite, G. A. O’Toole

**Author notes:** To whom correspondence should be addressed, Department of Microbiology and Immunology, Geisel School of Medicine at Dartmouth, Rm. 202 Remsen Building, 66 North College Street, Hanover, NH, 03755.

## Abstract

*Streptococcus sanguinis* is a prevalent member of human microbiome capable of acting as a causative agent of oral and respiratory infections. *S. sanguinis* competitive success within the infection niche is dependent on acquisition of metal ions and vitamins. Among the systems that bacteria use for micronutrient uptake is the energy coupling factor (ECF) transporter system EcfAAT. Here we describe physiological changes arising from EcfAAT transporter disruption. We found that EcfAAT contributes to *S. sanguinis* antibiotic sensitivity as well as metal and membrane homeostasis. Specifically, our work found that disruption of EcfAAT results in increased polymyxin susceptibility. We performed assessment of cell-associated metal content and found depletion of iron, magnesium, and manganese. Furthermore, membrane composition analysis revealed significant enrichment in unsaturated fatty acid species resulting in increased membrane fluidity. Our results demonstrate how disruption of a single EcfAAT transporter can have broad consequences on bacterial cell homeostasis. ECF transporters are of interest within the context of infection biology in bacterial species other than streptococci, hence work described here will further the understanding of how micronutrient uptake systems contribute to bacterial pathogenesis.

**Importance:** Proficiency in micronutrient uptake is key for pathogen success in bacteria-bacteria and bacteria-host interactions within the infection context. Micronutrient uptake mechanisms are of interest in furthering the understanding of bacterial physiology within infection niche and as targets for design of antimicrobials. Here we describe how a deletion of a nutrient uptake transporter in *S. sanguinis* alters bacterial sensitivity to antibiotics. We also show that a defect in this candidate nutrient uptake system has consequences on the intracellular metal content, and also results in changes in membrane fatty acid composition and fluidity. This study demonstrates how disruption of a single nutrient uptake system disrupts bacterial physiology resulting in increased antibiotic sensitivity.

## Introduction

*Streptococcus* is a diverse genus of Gram-positive bacteria whose species are both part of healthy human microbiome and capable of causing disease. *Streptococcus sanguinis* is commonly known to colonize oral cavity, where it’s presence is increased in association with disease (1). Additionally, streptococci are of increased interest in the context of lower airway infections and endocarditis (2). This organism is of special interest in the context of cystic fibrosis (CF) - a multiorgan genetic disease that is associated with chronic lung infections (3).

*S. sanguinis* infection physiology has been previously studied in context of bacteria-bacteria interactions occurring both in oral cavity and lungs (1, 4–6). Multiple studies implicate metal uptake as a key factor in *S. sanguinis* fitness within bacteria-bacteria competition. A screen examining *S. sanguinis* survival in presence of *Pseudomonas aeruginosa* (5) and an independent screen of *S. sanguinis* growth in nutritional conditions modeling lung infection found that deletions of any the genes in the three gene operon SSA2365-SSA2367 (6) result in a growth defect. Mutations in this same gene cluster were also found to significantly impact *S. sanguinis* growth in presence of human serum (7). Sequence based functional and structural domain prediction annotates the genes within this operon as encoding components of an energy coupling factor (ECF) transporter.

ECF transporters are a subclass of the adenosine 5’-triphosphate (ATP)-binding cassette (ABC) transporter superfamily. Unlike most ABC transporters that are present across prokaryotes and eukaryotes, ECF transporters have only been found encoded in prokaryotic genomes (8, 9). ECF transporters are comprised of two nucleotide binding domain-containing proteins termed EcfA and EcfA’, and a membrane integral protein - EcfT. EcfA and EcfT components comprise an energy coupling complex that interacts with a substrate binding proteins called the “S component” (8, 9). Individual EcfA-EcfT assemblies can interact with multiple substrate binding proteins that can be encoded in adjacent or remote genomic locations (10). Genes SSA2366 and SSA2367 are homologous to ECF A components, accordingly named EcfA2 and EcfA1, while SSA2365 is the transmembrane component termed EcfT. The genomic organization of SSA2365-67 gene cluster is consistent with these genes encoding a group II ECF transporter (9) where ATPase and transmembrane subunits are encoded in a single operon without an adjacent candidate gene for substrate binding protein (**Figure 1A, Suppl Table 1**). The ECF core components, lacking a substrate binding protein, that are encoded in the *S. sanguinis* SK36 SSA2365-67 cluster, are referred to here as EcfAAT.

**Figure 1.**
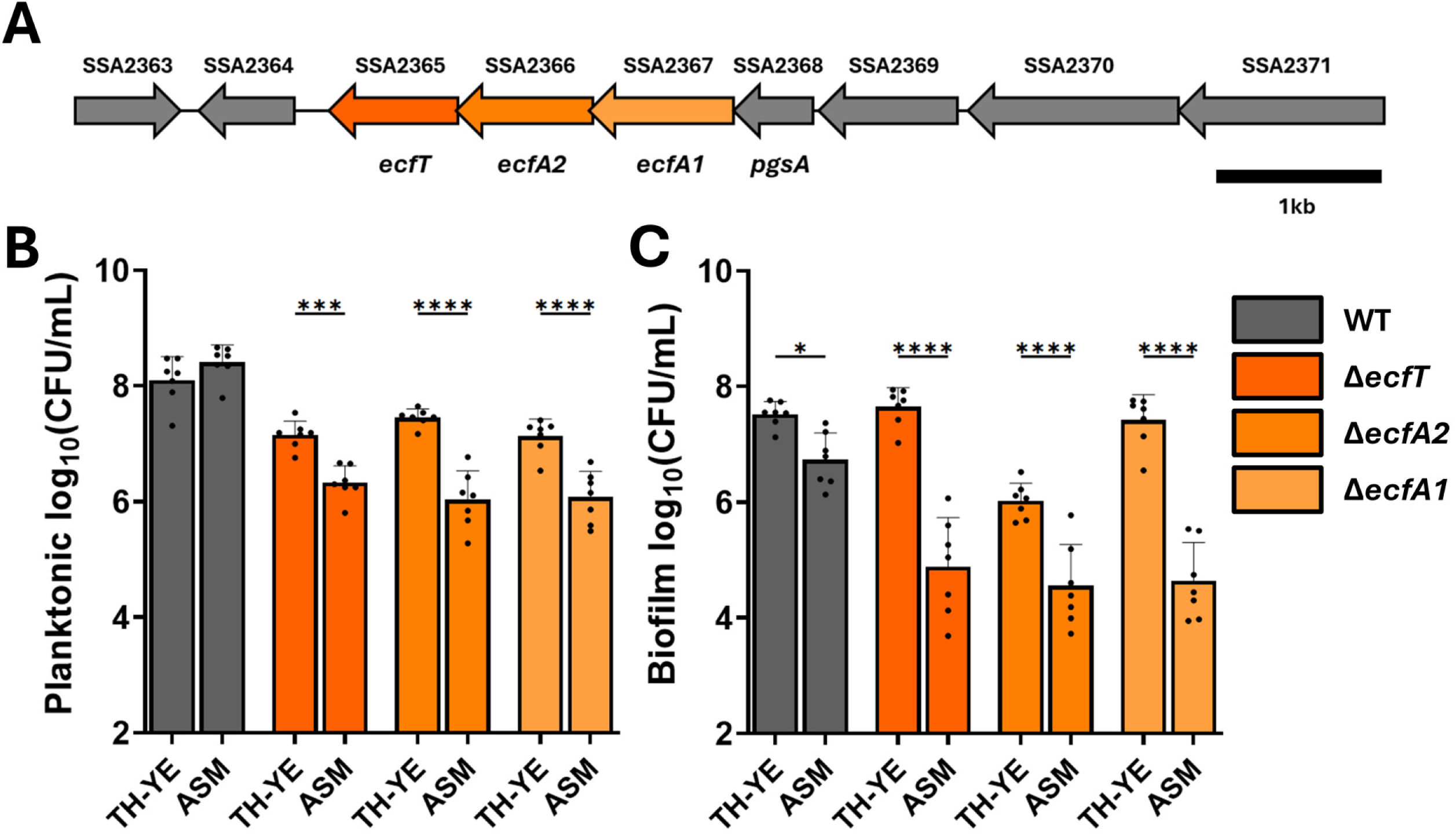
Artificial sputum media limits *ecfAAT* mutant growth and biofilm formation. (**A**) Schematic of *S. sanguinis* SK36 *ecfT-ecfA2-ecfA1* gene cluster organization, flanking gene descriptions detailed in **Supplementary Table 1**. Planktonic growth (**B**) and biofilm formation (**C**) of the WT and *ecfAAT* mutants compared in Todd-Hewitt broth with yeast extract (TH-YE) and artificial sputum medium (ASM). CFU counts assessed after 6h of static growth under anoxic conditions. Mean and standard deviation of n=7 biological replicates. Statistical analysis using ANOVA with Sidak’s post hoc test, with *, p<0.05, ***, p<0.001, ****, p<0.0001.

ECF transporters act strictly in uptake of small molecules, with specificity for compounds that are used in small quantities including enzymatic cofactors, such as vitamins or divalent cations (10, 11). ECF transporters in group A streptococci and *Staphylococcus lugdunensis* have been shown to contribute to uptake of heme and promote infection (12, 13). While another isolate of group A streptococcus was found to utilize horizontally acquired ECF S component for folate uptake leading to sulfamethoxazole resistance (14). ECF transporters can be found across prokaryotic genera with specific enrichment in the firmicutes (10). These features have highlighted ECF transporters as a novel target of interest in design of antimicrobial agents (15–17).

Given that ECF transporters are of emerging interest in context of *Streptococcus spp.* infection biology, we assessed how disruption of this transporter impacts antibiotic susceptibility. Our results show that strains lacking functional EcfAAT are more sensitive to polymyxin class antibiotics. To gain an understanding of the physiological changes induced by EcfAAT component deletion, we analyzed changes in the cell-associated metal content and have identified multiple putative EcfAAT substrates. Furthermore, we analyzed changes in *ecfAAT* mutant membrane composition and found that strains with an ECF transporter defect have increased membrane fluidity and are enriched in unsaturated fatty acid species. These data bring novel insights into the downstream effects of EcfAAT disruption, which will provide useful mechanistic information for studies aimed at designing antimicrobials targeting ECF transporters.

## Results

### ECF transporter loss results in growth and biofilm formation defect

*S. sanguinis* strains with energy coupling factor (ECF) transporter gene deletions have been previously found to have growth defect when exposed to infection niche-relevant conditions (5, 7). In this study, we evaluated fitness of strains lacking genes encoding individual EcfAAT components when grown in nutritional conditions mimicking CF sputum (artificial sputum medium, ASM) while under anoxic atmosphere that best reflect conditions within lung infection environment (18, 19). We assessed fitness of mutants lacking individual *ecfA1*, *ecfA2*, and *ecfT* genes and observed that these strains have a significant impairment in both planktonic and surface-attached growth as compared to the WT strain when grown in undefined laboratory medium conditions (Todd-Hewitt broth supplemented with yeast extract, TH-YE; **Figure 1B and C**). The EcfAAT mutant growth defect was further exaggerated when strains were cultured in ASM (**Figure 1B and C**).

By growing *S. sanguinis* mutants in a 1:1 mixture of rich laboratory medium (TH-YE) and ASM, we observed that *ecfAAT* mutant growth as a biofilm was significantly higher in medium containing Todd-Hewitt broth, indicative of ASM lacking one or more nutrients required for biofilm establishment by the *ecfAAT* mutants (**Suppl Figure 1**). In addition to the numerical growth defect quantified, the *ecfAAT* mutant strains colonies are consistently smaller in size (not shown). To confirm that the observed growth defect is a result of the specific gene deletions, we reintroduced the missing genes into an ectopic site of *S. sanguinis* genome. Using *ecfAAT* mutant complementation strains, we saw that the restoration of the missing gene enabled strains to grow to the same extent as WT (**Suppl Figure 2**).

### Loss of EcfAAT transporter results in decreased intracellular iron, manganese, and magnesium

ECF transporters have been described to act strictly as importers involved in uptake of small molecules that typically function as co-factors or co-factor precursors (10). Substrates identified to date include divalent cations, amino acids, and vitamins such as biotin, folate, riboflavin, or cobalamin (8, 10). KEGG functional prediction classified *S. sanguinis* SK36 EcfAAT as a transporter associated with iron-siderophore, cobalt, and vitamin B_12_ metabolism.

To investigate the potential substrates of EcfAAT, we used inductively coupled plasma mass spectrometry (ICP-MS) to assess changes in cell associated metal content. This analysis measured concentration of a 16-metal panel (**Figure 2, Suppl Figure 3, and Suppl Table 2**) of washed bacterial cell pellets adjusted to the weight of the pellet. We found that all three *ecfAAT* mutants have an average of 40-50% less intracellular iron (Fe_WT_=36±7.4ng/mg; Fe_EcfT_=22.3±5.2ng/mg; Fe_EcfA2_=23±2.7ng/mg; Fe_EcfA1_=20.3±4.8ng/mg) and manganese (Mn_WT_=47.1±8.8ng/mg; Mn_EcfT_=23±2.1ng/mg; Mn_EcfA2_=22.2±3.3ng/mg; Mn_EcfA1_=23.7±5.2ng/mg) compared to WT *S. sanguinis*. Additionally, cell-associated magnesium levels are also significantly decreased (Mg_WT_=1.3±0.06µg/mg; Mg_EcfT_=1.08±0.15µg/mg; Mg_EcfA2_=1.06±0.04µg/mg; Mg_EcfA1_=0.97±0.03µg/mg). Although functional domain conservation analysis predictions associate EcfAAT components with cobalt uptake, we saw no significant changes in the amounts of cell-associated cobalt (Co_WT_=8.2±1.3ng/mg; Co_EcfT_=8.5±1.2ng/mg; Co_EcfA2_=8.9±1.7ng/mg; Co_EcfA1_=8.8±1.7ng/mg). Similarly, our analysis did not detect significant changes in the zinc (Zn_WT_=54±5.2ng/mg; Zn_EcfT_=52.4±2.7ng/mg; Zn_EcfA2_=54.1±0.8ng/mg; Zn_EcfA1_=52.6±1.5ng/mg), or calcium content (Ca_WT_=40±1.8ng/mg; Ca_EcfT_=30.1±7.8ng/mg; Ca_EcfA2_=46.4±20.1ng/mg; Ca_EcfA1_=32.2±7.2ng/mg). The full trace element panel (**Suppl Figure 3, and Suppl Table 2**) showed a consistent decrease in the mean cadmium concentration, with the Δ*ecfA1* mutant being significantly different from WT (Cd_WT_=0.32±0.06ng/mg; Cd_EcfA2_=0.2±0.05ng/mg). Additionally, strontium measurements showed a significant decrease in both Δ*ecfT* and Δ*ecfA1* mutants (Sr_WT_=0.1±0.01ng/mg; Sr_EcfT_=0.05±0.02ng/mg; Sr_EcfA1_=0.06±0.01ng/mg). This metal content analysis has identified multiple putative EcfAAT substrates. However, further analysis would be needed to assess weather changes in the metal content are direct result of impairment in the specific metal uptake or general disruption in metal metabolism.

**Figure 2.**
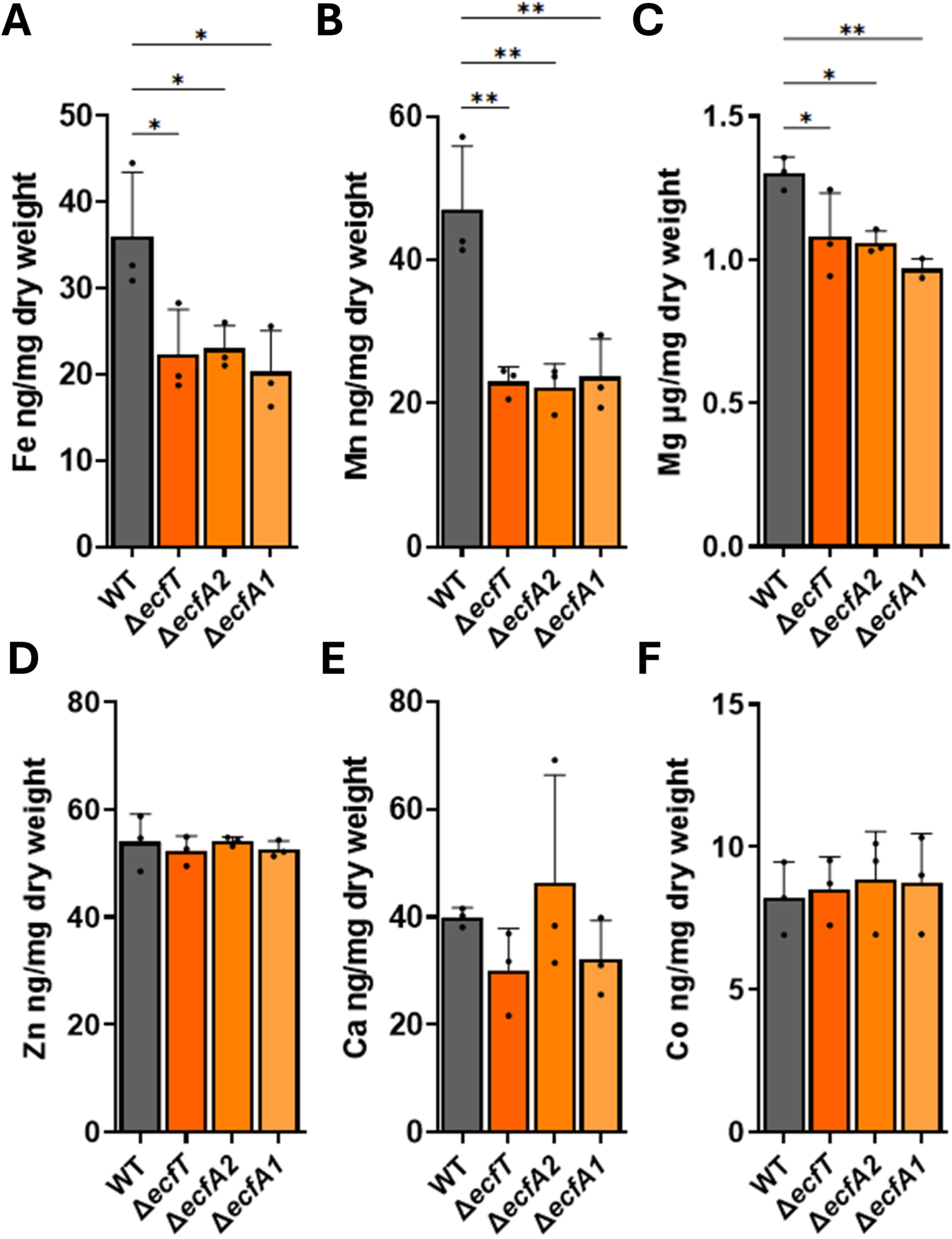
The EcfAAT transporter contributes to metal uptake. WT and mutant cell-associated metal content assessed by ICP-MS using bacteria grown in TH-YE media for 6h at 37°C, 5% CO_2_. Cell associated content of iron (**A**), manganese (**B**), magnesium (**C**), zinc (**D**), calcium (**E**), and cobalt (**F**). The values for the 10 additional metals assessed are shown in **Supplementary Figure 3** and **Supplementary Table 2**. Metal content reported as ng or µg per mg of dry cell weight. Mean and standard deviation of n=3 biological replicates shown. Statistical analysis using ANOVA with Dunnett’s post hoc test, with *, p<0.05, **, p<0.01.

### ECF mutants show increased sensitivity to polymyxin antibiotics

As the *ecfAAT* mutant growth defect is exaggerated ASM compared to growth in nutritionally undefined laboratory media conditions (**Figure 1, Suppl Figure 1**), we aimed to further evaluate clinically-relevant impact of these mutations. For these studies, we assessed whether loss of EcfAAT components affects antibiotic sensitivity. We observed no consistent differences in susceptibility to Vancomycin, Clindamycin, Ciprofloxacin or Levofloxacin (**Suppl Figure 4A-D**). In contrast, all of the *ecfAAT* mutants display increased sensitivity to polymyxin class antibiotics – colistin (Polymyxin E) and Polymyxin B (**Figure 3**). While WT MIC for Polymyxin B is 512µg/mL, the mutant strains are sensitive to 256µg /mL, and corresponding colistin MICs are 1024µg/mL and 512µg/mL, respectively.

**Figure 3.**
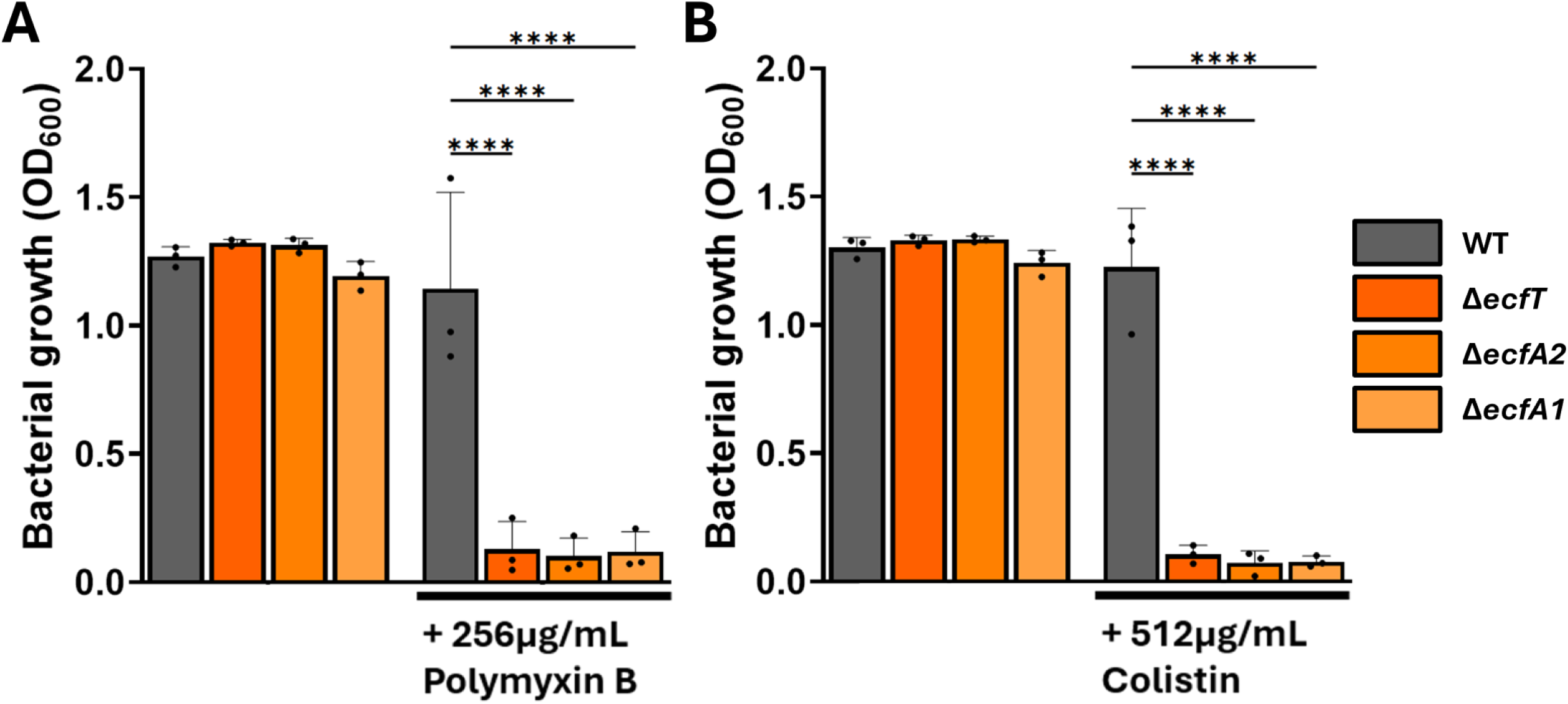
EcfAAT transporter defect leads to increased polymyxin sensitivity. To assay the impact of mutating the EcfAAT system on antibiotic sensitivity we exposed WT and *ecfAAT* mutant set to Polymyxin B (**A**) or colistin (**B**) treatment. For this assay, bacteria were inoculated into TH-YE medium containing the specified amount of a given antibiotic. Bacterial growth was assessed by OD_600_ measurements following 18h static growth at 37°C, 5% CO_2_. Mean and standard deviation of n=3 biological replicates shown. Statistical analysis using ANOVA with Tukey’s post hoc test, with, ****, p<0.0001.

Polymyxin antibiotics act by disrupting bacterial cell wall and membrane integrity (20, 21). To address whether the *ecfAAT* mutants are sensitive to polymyxins specifically or whether these strains show a more general increase in sensitivity to membrane and cell wall targeting antibiotics we next assessed changes in sensitivity to daptomycin. In our assay conditions, we saw no consistent change in sensitivity to daptomycin when comparing *S. sanguinis* SK36 WT and any of the *ecfAAT* mutants, with an MIC of 32µg/mL for all these strains (**Suppl Figure 4E**).

### Ca and Mg protect *S. sanguinis* from Polymyxin B toxicity

Polymyxin molecular targets in both Gram-positive and -negative bacteria are LPS or membrane domains rich in negative charge (22–25). These structures are stabilized by divalent cations such as Ca^2+^ or Mg^2+^, and polymyxin interactions with cellular targets are reliant on displacement of these ions (26–28). Cation supplementation has been shown to be protective against polymyxin toxicity (29, 30). As changes in cation homeostasis have been shown to impact polymyxin sensitivity in other bacterial species, we evaluated whether differences in the cell associated metal concentrations could account for the increased sensitivity to Polymyxin B of the EcfAAT mutants.

To investigate how addition of metal ions impacts EcfAAT mutant antibiotic susceptibility we employed a checkerboard assay. First, we tested impact of Mg addition, as this metal shows the highest magnitude of depletion for the *ecfAAT* mutant cells compared to the WT (**Figure 2**). Supplementation of 10mM of Mg^2+^ appears to consistently restore the Polymyxin B sensitivity levels of the Δ*ecfT* mutant to nearly WT levels, with growth being detectable in the presence of 512µg/mL of Polymyxin B (**Figure 4**). Notably, high concentrations of added MgCl_2_ are also protective of WT *S. sanguinis* enabling growth in presence of 1024µg/mL of Polymyxin B.

**Figure 4.**
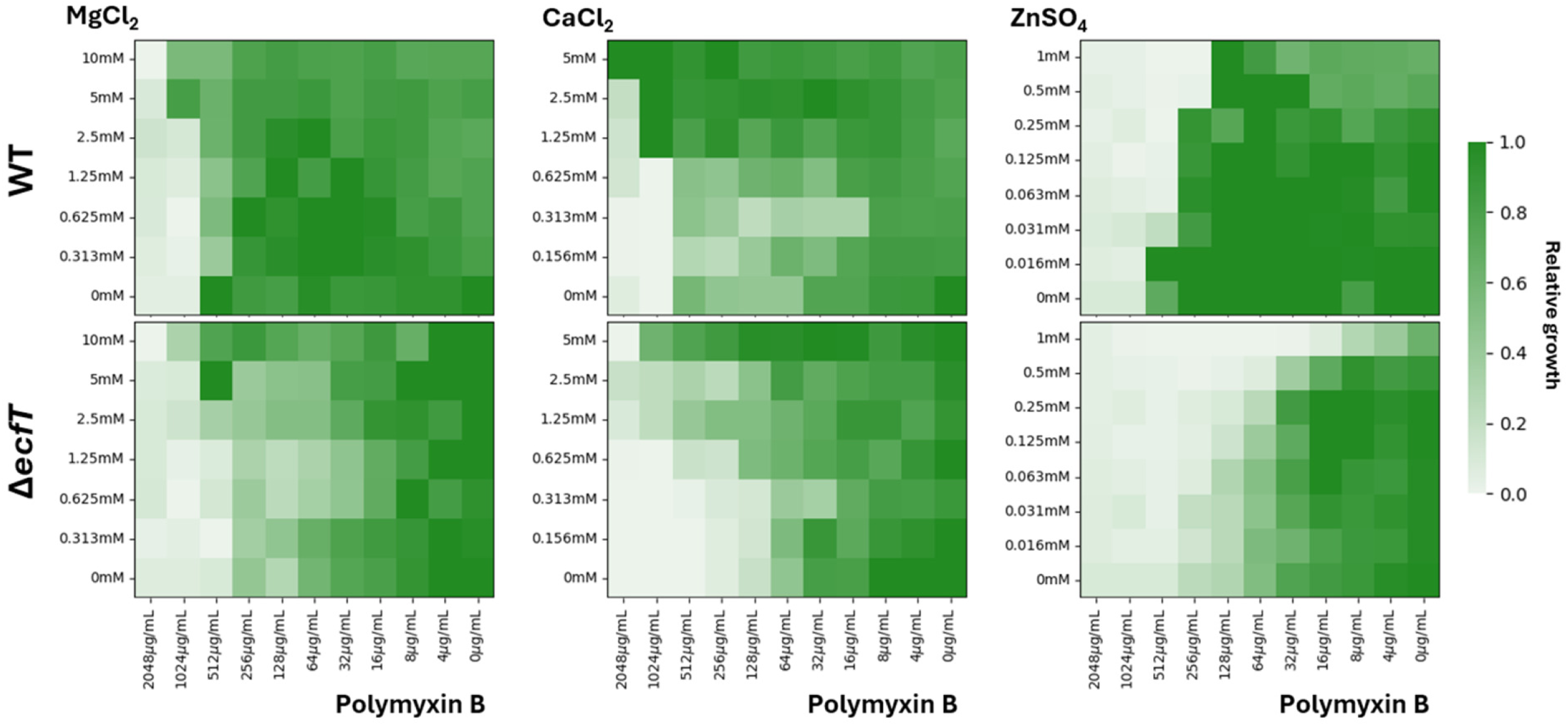
Addition of metal ions alters Polymyxin B antimicrobial activity. Checkerboard assay assessing how combined metal ion and Polymyxin B exposure impacts WT and Δ*ecfT* strain growth. Both magnesium (0.3125mM to 10mM) (**Panels on left**) and calcium (0.15625mM to 5mM) (**Central panels**) addition protects *S. sanguinis* from Polymyxin B toxicity in a dose dependent manner. Supplementation of zinc (0.015625mM to 1mM) (**Panels on right**) increases Polymyxin B toxicity. Measurements performed by inoculating WT (**Top row**) or Δ*ecfT* mutant (**Bottom row**) into TH-YE media containing a metal and antibiotic mixture. Optical density measurements were performed after 18h static growth at 37°C, 5% CO_2_ atmosphere. A representative measurement set of 3 biological replicates shown, with darker shading indicating higher bacterial amount.

To assess whether this protective effect extends beyond cations depleted in the *ecfAAT* mutants, we tested impact of calcium supplementation. Addition of high concentrations of CaCl_2_ enabled WT *S. sanguinis* growth in presence of 2048µg/mL of Polymyxin B and growth of the Δ*ecfT* mutant in the presence of 1024µg/mL of Polymyxin B. The observed effect of both Ca^2+^ and Mg^2+^ ions protecting WT cells from Polymyxin B indicates a more general protective mechanism than restoration of the ions depleted in the *ecfAAT* mutants.

The above-described checkerboard assays were performed by inoculating bacteria directly into media containing antibiotics. Although this is a common approach for MIC testing, this method assesses the sensitivity of planktonic cells to antibiotics. However, previous research has shown that bacteria within the infection niche often exist in a form of a biofilm (Costerton, Stewart, and Greenberg 1999; Braxton et al. 2005). To test whether addition of metal ions protects established biofilms from Polymyxin B toxicity, we next adapted the above checkerboard assay to assess established biofilm antibiotic treatment tolerance. Here we allowed for *S. sanguinis* biofilm establishment for 18h, before exposure to the metals and antibiotic treatment mixtures. The assay data showed that the Δ*ecfT* mutant biofilms remained more sensitive to Polymyxin B than WT (**Suppl Figure 5**). Our assay is not able to distinguish whether this is a result of inherent higher sensitivity of the strains and/or comparable poor biofilm establishment prior to treatment exposure resulting in numerically smaller starting population compared to WT. Overall, these assays reflected Ca^2+^ and Mg^2+^ ions acting antagonistically with Polymyxin B in the same manner as described above for planktonic bacteria.

Finally, optical density readings were indicative of partial bacterial growth in the presence of even the highest antibiotic concentrations without Ca^2+^ or Mg^2+^ addition. To investigate whether viable bacteria are present or measurements are reporting biofilm debris, we cultured the remaining biomass onto non-selective medium to allow for viable bacteria recovery. Little to no bacteria were detected when plating contents of these wells, indicative of Polymyxin B having a bactericidal effect that allowed for killing of the bacteria within the established biofilm rather than simply inhibiting further growth.

### High zinc concentrations act synergistically with Polymyxin B

While Ca^2+^ and Mg^2+^ ions have been reported to act by stabilizing bacterial membrane and cell wall, a previous investigation demonstrated that addition of ionophore PBT2 results in increased sensitivity to Polymyxin B in a Zn^2+^ dependent manner (31). Zinc is of interest within the context of infection niche as it is highly abundant in CF, essential for bacterial survival, and involved in host-bacteria and bacteria-bacteria interactions (5, 32, 33). Using the checkerboard assay, we saw that ZnSO_4_ synergizes with Polymyxin B with combined treatment enhancing antimicrobial activity versus the Δ*ecfT* mutant (**Figure 4**). Addition of 0.5mM or 1mM of Zn^2+^ shifted WT Polymyxin sensitivity from 512µg/mL to 256µg/mL, with even further increase in Polymyxin B sensitivity demonstrated by the Δ*ecfT* mutant (1mM ZnSO_4_ addition resulted in growth eradication at 16-32µg/mL). ZnSO_4_-Polymyxin B synergy observation was also confirmed when assessing biofilm-grown bacteria (**Suppl Figure 5**), although the effect versus biofilm-grown bacteria was more modest.

### Loss of EcfAAT does not result in measurable cell wall defect

The general polymyxin ineffectiveness against Gram-positive bacteria is largely due to the physical barrier provided by the peptidoglycan layer, as both *S. aureus* and *Bacillus subtilis* protoplasts are sensitive to Polymyxin B treatment (34, 35). To investigate whether *ecfAAT* mutant sensitivity to polymyxins is a result of cell wall defect we imaged WT and mutant bacteria using transmission electron microscopy (TEM). Using this methodology, we were unable to detect any consistent defects in cell wall morphology (**Suppl Figure 6A**). Additionally, our measurements did not show significant changes in the mean cell wall thickness when comparing WT and *ecfAAT* mutant cells (**Suppl Figure 6B**).

Teichoic acids (TAs) and lipoteichoic acids (LTAs) are anionic glycopolymers present in Gram-positive bacteria cell wall (36, 37). Polymyxin molecules have been demonstrated to interact with TAs (25). Additionally, charge reducing modifications of TAs occur in a range of Gram-positive organisms and have been shown to contribute to polymyxin resistance of *Bacillus thuringiensis* (38, 39). A similar protective effect is seen in cases of charge reducing aminoacylation of phospholipid headgroups (39, 40). To address whether increase in Polymyxin B sensitivity observed for the *ecfAAT* mutants is a result of an overall change in the cell surface charge we performed zeta potential measurements of the strains of interest. To obtain zeta potential measurements bacteria are placed in an electrophoresis capillary and differences in cell migration are related to an overall change in the surface charge (41, 42). Our zeta potential measurements did not detect significant changes in the overall cell surface charge when comparing WT and the *ecfAAT* mutant cells (**Suppl Figure 6C**).

### Loss of EcfAAT leads to increased membrane fluidity

Another aspect of bacterial cell physiology described to impact polymyxin sensitivity is changes in membrane integrity (20, 43). To address whether mutations of the genes coding for the EcfAAT transporter have an impact on membrane integrity we utilized the Laurdan general polarization (GP) assay. These measurements rely on a membrane integral fluorophore shifting light emission wavelength depending on the water content within membrane. These shifts in fluorescence are sensitive to changes in phospholipid head group density and fatty acyl spreading – jointly describing changes in membrane fluidity (44–46).

The *ecfAAT* mutant strains show a significant reduction in Laurdan GP compared to WT, indicative of relative increase in membrane fluidity (**Figure 5A**). This observation is consistent with the mutant strains increased polymyxin sensitivity, as reduction in phospholipid packing would allow for increased polymyxin integration into bacterial membranes (47). Exposure to Polymyxin B leads to a significant increase in Laurdan GP in both WT and *ecfAAT* mutants (**Figure 5B** and **Suppl Figure 7**). The observed polymyxin induced increase in membrane rigidity is consistent with previous observations for *E. coli* (23, 48).

**Figure 5.**
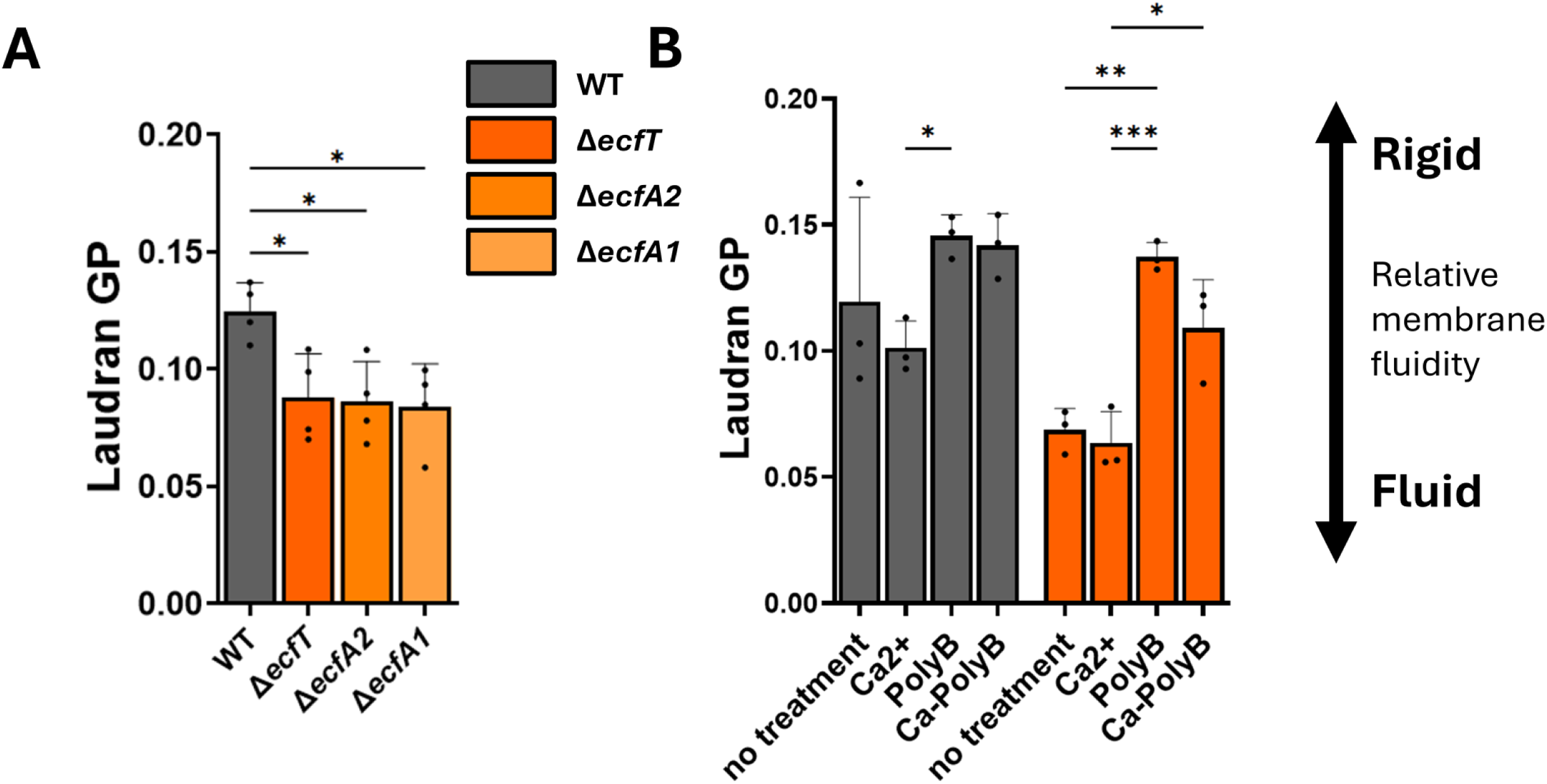
*S. sanguinis* membrane fluidity is influenced by mutations in the EcfAAT transporter. As measured using Laurdan generalized polarization (GP) assay, (**A**) *S. sanguinis* strains with mutations in the EcfAAT-encoding genes have a significantly less rigid cell membrane compared to WT. (**B**) Impact of individual and combined Ca^2+^ and Polymyxin B treatment on WT and Δ*ecfT* mutant membrane fluidity. Before analysis, bacteria were cultured statically, in TH-YE medium, at 37°C, 5% CO_2_. Mean and standard deviation of n=5 biological replicates shown. Statistical analysis using ANOVA with Dunnett’s post hoc test.

The measurements described in **Figure 3** show that addition of metal ions impacts polymyxin effectiveness, therefore we tested weather addition of Ca^2+^ or Zn^2+^ has an impact on membrane fluidity that could explain changes in the polymyxin susceptibility. Addition of high concentrations of Ca^2+^ or Zn^2+^ ions did not result in significant changes in WT or mutant strain membrane fluidity (**Figure 5B** and **Suppl Figure 7**). Notably, although combined addition of Polymyxin B and Ca^2+^ still resulted in significant elevation in membrane rigidity, this change occurred to a lesser extent than treatment with only Polymyxin B (**Figure 5B**), while Zn^2+^ addition did not affect Laurdan GP regardless of Polymyxin B addition (**Suppl Figure 7**).

### The impact of loss of EcfAAT function on membrane composition

To address weather shifts in membrane fluidity displayed by *ecfAAT* mutants are a result of changes in the overall membrane composition, we submitted the WT and mutant strains to fatty acyl methyl ester (FAME) analysis. This analysis included 24 FAME species with abundance of more than half of these being significantly shifted in *ecfAAT* mutants (**Figure 6, Suppl Figure 8**, and **Suppl Table 3**). Membrane composition of all three of *ecfAAT* mutants was shifted in the same manner, compared to WT. The overall fraction of saturated FAME species was decreased by approximately 50% in mutant strains compared to the WT, and correspondingly both mono- and poly-unsaturated FAME species were more prevalent.

**Figure 6.**
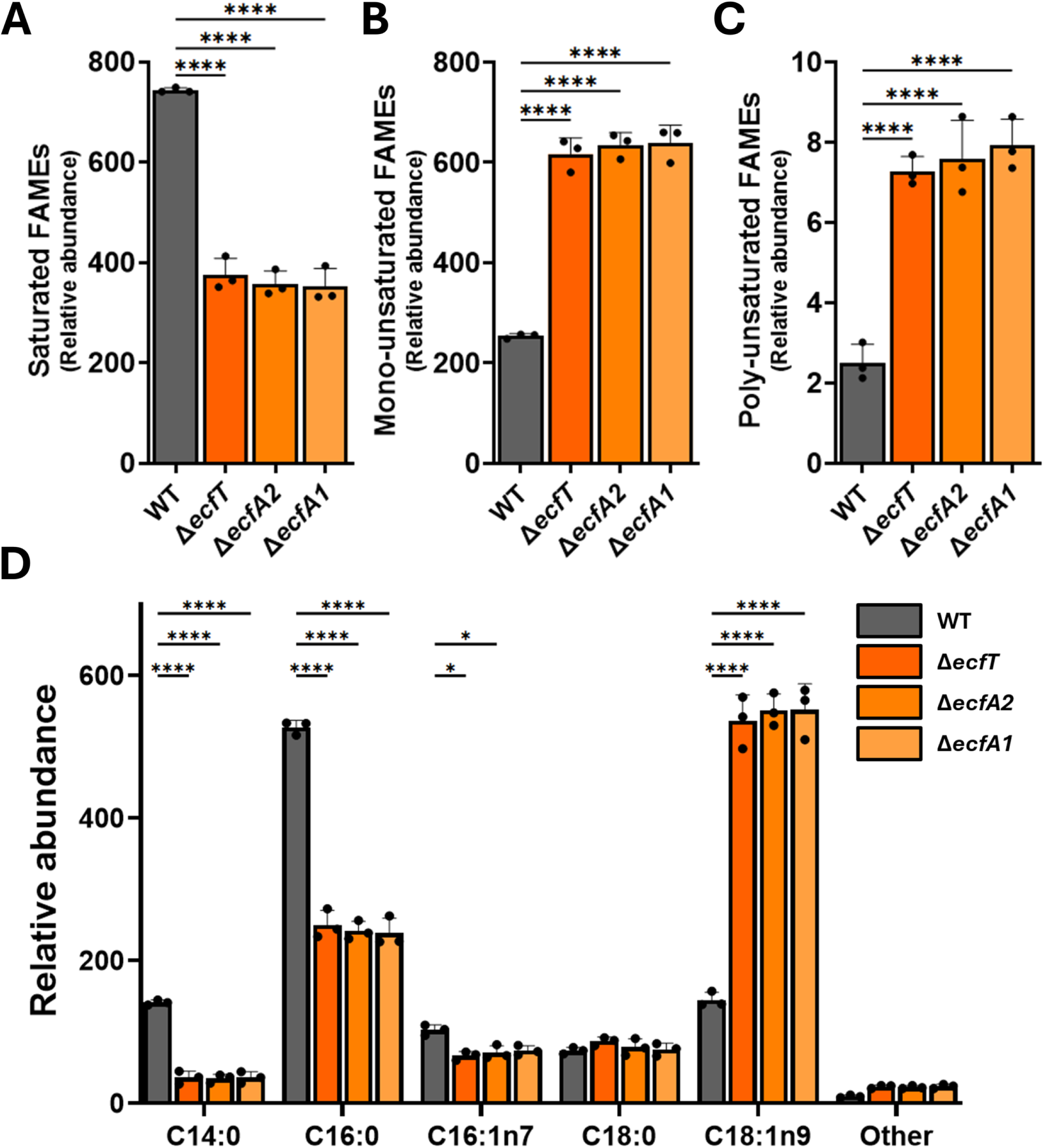
*S. sanguinis* strains with EcfAAT transporter defect have significantly altered membrane composition. To assess changes in bacterial membrane composition, WT and *ecfAAT* mutant strains were subjected to fatty acyl methyl ester (FAME) analysis. Summary of relative changes in saturated (**A**), mono-unsaturated (**B**), and poly-unsaturated (**C**) FAME content in WT and *ecfAAT* mutant strains. (**D**) Changes in the five most abundant FAME content – C14:0; C16:0; C16:1n7; C18:0; C18:1n9; with “other” category summing additional 19 FAME species analyzed. Full FAME analysis panel results are detailed in **Suppl Figure 8** and **Suppl Table 3**. For the purposes of this analysis, bacteria were cultured statically, in TH-YE media, at 37°C, 5% CO_2_. Mean and standard deviation of n=3 biological replicates shown. Values shown as adjusted to the total FAME content. Statistical analysis using ANOVA with Dunnett’s post hoc test.

This shift was accounted for by a substantial depletion of myristic (C14:0) and palmitic (C16:00) fatty acids, in favor of increased oleic (C18:1n9) fatty acid content. Both, major and minor FAME species analysis showed increased relative abundance of longer FAME species. The enrichment in unsaturated FAME species and increase in the overall chain length is consistent with the above observed increase in bacterial membrane fluidity.

## Discussion

Our work shows that disruption of *S. sanguinis* EcfAAT transporter homolog impacts cellular metal homeostasis and membrane integrity resulting in increased antibiotic susceptibility. Disruption of the EcfAAT transporter has been previously described to result in a growth defect in presence of serum (7). Additionally, previous screens have found deletions of this gene to result in growth impairment in artificial sputum medium (6), as well as impact *S. sanguinis* and *P. aeruginosa* interactions in co-culture (5). These observations here highlight EcfAAT as a molecular target of interest in context of *S. sanguinis* pathogenesis as it’s disruption has implications for strain fitness under growth conditions similar to those found in the CF lung. Here we describe that deletion of the genes coding for any component of the putative EcfAAT transporter results in not only cell-associated metal depletion but also significantly alters bacterial membrane composition and fluidity.

Common ECF transporter substrates include vitamins and metal ions (10, 11) and homology-based functional predictions assign *S. sanguinis* EcfAAT transporter as contributing to cobalt or cobalamin uptake. Our analysis (**Figure 2**) did not detect changes in cell associated cobalt concentrations, a finding in an agreement with a prior analysis by (7). A miss-annotation classifying ECF transporter components as belonging to cobalt (Cbi) or nickel (Nik) uptake systems has been reported previously (10). Our cell associated metal content analysis detected significant changes in iron, manganese, magnesium, cadmium, and strontium levels (**Figure 2**, **Suppl Figure 3**, and **Suppl Table 2**). These metal ions are a set of potential EcfAAT substrates, however confirmatory work would be reliant on identification of the specific substrate binding components associated with the EcfAAT transporter. Identifying such binding components would subsequently allow us to pinpoint which of these are direct EcfAAT substrates and which metal levels may be disrupted indirectly. As EcfAAT is a predicted type II ECF transporter capable of associating with multiple distinct substrate binding components, it is possible that EcfAAT substrate set could also include other small molecules such as vitamins.

Antimicrobial susceptibility testing of the *ecfAAT* mutants revealed a modest increased susceptibility to polymyxin class antibiotics (**Figure 3**). Polymyxins are positively charged cyclic lipopeptide antibiotics that induce membrane damage (43, 49). Polymyxins preferentially interact with the negatively charged phospholipids, lipopolysaccharides (LPS), and lipid A specifically (20, 50). Lipid A target specificity is the reason for polymyxins being considered largely ineffective against Gram-positive bacteria including most streptococci (39). However, polymyxins can disrupt membranes of Gram-positive protoplasts (34, 35) indicating that protection is provided by the Gram-positive cell wall. Our analysis did not detect changes in *S. sanguinis* cell wall thickness or overall surface charge (**Suppl Figure 6**). Polymyxin mechanism of action in Gram-negative bacteria is dependent on displacement of cell wall and cell membrane associated calcium and magnesium ions (51, 52). Subsequently, supplementation with these metals has been reported to protect *P. aeruginosa*, *Acinetobacter* spp. and other microorganisms from polymyxin toxicity (53–55). Our analysis revealed that high concentrations of Ca^2+^ or Mg^2+^ ions protect *S. sanguinis* from Polymyxin B toxicity (**Figure 4** and **Suppl Figure 5**). Although *S. sanguinis* lacks lipid A that acts as polymyxin molecular target in Gram-negative bacteria, it appears that cation-mediated stabilization of the cell wall and membrane (56) could still be an important physiological factor contributing to polymyxin tolerance.

Changes in membrane composition have been reported to impact bacterial susceptibility to polymyxins (21). Our measurements revealed that *ecfAAT* mutant strains displayed increased relative membrane fluidity compared to WT *S. sanguinis* (**Figure 5**), while addition of Polymyxin B resulted in increased membrane rigidity that was in part inhibited by addition of Ca^2+^. These observations are consistent with a previously proposed model where polymyxin molecules have to compete with Ca^2+^ ions when interacting with bacterial cell wall and membrane and high Ca^2+^ ion concentration can act to prevent polymyxin integration into bacterial cell membrane consequently decreasing toxicity (30, 57). Subsequent membrane composition analysis revealed that mutant strain membranes are enriched in unsaturated fatty acids and fatty acids with longer chain length (**Figure 6**, **Suppl Figure 8**, and **Suppl Table 3**). These changes in the membrane fatty acid content are consistent with the observed increase in membrane fluidity.

Further investigation would be required to address the mechanistic reasons leading to these changes in the membrane fatty acid content composition. Here we propose three potential directions for future investigation of this effect. First, these changes in the membrane could be a direct result of the depletion of the EcfAAT transporter substrates, wherein changes in membrane composition are a result of a compensatory mechanism in response to ion and other micronutrient depletion. Secondly, as EcfAAT substrates are common enzyme co-factors, loss of these co-factors could indirectly impact membrane biosynthesis. A previous study investigated *S. aureus* small colony variant mutants, which were found to have an ECF transporter defect. These strains showed auxotrophy for unsaturated fatty acids, and authors describe overall phenotypic similarities with vitamin uptake auxotrophic strains of *S. aureus* (58). A third point of consideration is genomic location of EcfAAT operon (**Figure 1**); these genes are encoded downstream of an essential phospholipid synthesis enzyme - phosphatidylglycerol phosphate (PGP) synthase (PgsA) (59). PgsA defects have been shown to impact both phospholipid head group and fatty acid composition of streptococci membranes (60). Loss of PgsA, has been shown to lead to lead to increased membrane fluidity in *S. aureus*, but unlike in our work, this shift in the membrane leads to a high-level daptomycin resistance (61). We did not observe a change in daptomycin sensitivity in the mutants studies here. Further, our experimental work showed that complementation of EcfAAT components does restore bacterial growth to WT levels, but this does not fully exclude possibility of EcfAAT gene deletions affecting expression of an adjacent genomic locus.

ECF type transporters are bacterial-specific and broadly conserved, with enrichment in firmicutes (9). Therefore, study of these transporters is relevant not only to *Streptococcus sp*., but also other pathogens of interests for design of antimicrobial therapies including *Staphylococcus*, *Clostridium*, and *Enterococcus* species (15–17, 62). ECF transporters are involved in a range of micronutrient uptake. Furthermore, EcfAAT is a proposed type II ECF transporter that acts as a platform interacting with multiple distantly encoded substrate binding proteins (10, 11), so disruption of the EcfAAT functional unit may impact the uptake of a range of nutrients at once. These features position ECF transporters as excellent putative targets for novel antimicrobial therapy design (15). Multiple recent studies have reported screening of compounds targeting ECF transporters, including, *Lactobacillus* and *S. pneumonia* targeting compounds (16, 17, 62–64).

## Materials and methods

### Bacterial strains and growth conditions

Bacterial strains used in the study are listed in the **Supplementary Table 4**. *S. sanguinis* strains were routinely cultured on Tryptic soy agar plates supplemented with 5% v/v defibrillated sheep blood, or Todd-Hewitt (TH) broth supplemented with 0.5% w/v yeast extracts (TH-YE). When preparing overnight liquid cultures, a single colony was inoculated into a glass tube with 7mL of TH-YE, and the bacteria cultured at 37°C under 5% CO_2_ atmosphere without agitation. For purposes of microbial growth assays testing growth in different media, bacteria were grown under anoxic conditions in an anaerobic environmental chamber (Coy labs) with 5% CO_2_, 5% H_2_, and 90% N_2_ atmosphere without agitation. For purposes of MIC testing, membrane fluidity analysis, zeta potential analysis, and preparation of bacterial samples for mass spectrometry analysis cells were cultured under 5% CO_2_ atmosphere without agitation. *E. coli* strains were cultured in LB at 37°C with agitation. Spectinomycin was used at 50µg/mL for *E. coli* and 200µg/mL for *S. sanguinis* strains.

### Construction of *S. sanguinis* complementation plasmids

Gene complementation constructs were assembled using a suicide vector pJFP126 (65). Using this plasmid, genes are placed under an IPTG inducible promoter and inserted into the *S. sanguinis* chromosome at the site of the SSA0169 gene.

*S. sanguinis* SK36 genomic DNA was purified using DNeasy Blood & Tissue Kit, according to the manufacturer’s instructions for Gram-negative organisms. The *ecfT*, *ecfA2*, and *ecfA1* genes were individually amplified from the *S. sanguinis* genomic DNA using NEB Q5 High-Fidelity DNA Polymerase using primers specified in **Supplementary Table 5**. Primers were designed to amplify the entirety of the gene of interest and approximately 40 to 50 bp of the upstream promoter region. The amplified PCR fragments were purified using Qiagen QIAquick PCR purification kit and plasmid was purified using Qiagen QIAprep Spin Miniprep kit. Insert DNA and empty vector plasmids were digested using the following NEB enzymes according to the manufacturer’s instructions – *Hind*III, *Nhe*I, and *Sph*I. Subsequently, inserts were ligated into the plasmid backbone using NEB T4 Ligase according to the manufacturer’s instructions and chemically transformed into *E. coli* DH5α. Accuracy of plasmid construct (**Supplementary Table 6**) sequences was confirmed by sequencing at the Dartmouth Genomics and Molecular Biology Core.

### Transformation of *S. sanguinis*

*S. sanguinis* strains containing complementation plasmid inserts were constructed using a transformation protocol adapted from a previous report (66). Briefly, 50µL of *S. sanguinis* recipient strain overnight cultures were used to inoculate sub-culture into 10mL of fresh TH-YE media. After 3h growth at 37°C 5% CO_2_, 1mL of *S. sanguinis* subculture was supplemented with 100ng of competence stimulating peptide and mixed with 1µg of plasmid DNA. *S. sanguinis* SK36 competence stimulating peptide with the sequence of DLRGVPNPWGWIFGR was purchased from GenScript. Following incubation, *S. sanguinis* transformants were selected by growth on TSB agar supplemented with 5% v/v sheep’s blood and 200µg/mL of spectinomycin. Transformants containing complementation plasmid were screened using colony PCR using NEB Taq polymerase. Following the initial isolation of strains containing complementation constructs, strains were cultured without addition of antibiotics, and experiments were performed without IPTG induction, as initial testing showed that presence of the native promoter in combination with uninduced expression from hyper-spank promoter within the plasmid was sufficient to restore WT strain like phenotype within experimental conditions tested.

### Microbial growth assays

Bacterial growth in planktonic and biofilm fractions was assessed in a 96-well plate format. Bacteria from overnight cultures were aliquoted into microcentrifuge tubes, pelleted using a benchtop centrifuge (6000 x *g*, 3min) and subsequently washed in phosphate-buffered saline (PBS). After two wash steps, OD_600_ was measured, and bacterial culture densities were adjusted to an OD_600_=0.4. Subsequently, 50µL of bacteria were mixed with 950µL medium of interest, and 3 technical replicates of 100µL were transferred to a 96-well plate. Bacterial growth in 5 media conditions was evaluated: Todd-Hewitt broth supplemented with 0.5% yeast extract (TH-YE), TH-YE broth mixed with PBS in 1 to 1 ratio, artificial sputum medium (ASM), ASM mixed with PBS at a 1 to 1 ratio, and ASM mixed with TH-YE at a 1 to 1 ratio.

The ASM recipe used in this study was adapted from the SCFM2 recipe described previously (67) with modifications (68). Briefly, ASM with the following composition was used: Na_2_HPO_4_ (1.3mM), NaH_2_PO_4_ (1.25mM), KNO_3_ (0.348mM), K_2_SO_4_ (0.271mM), glucose (3mM), L-lactic acid (9.3mM), CaCl_2_ (1.754mM), MgCl_2_ (0.606mM), N-acetylglucosamine (0.3mM), tryptophan (0.066mM), 1,2-dioleoyl-sn-glycero-3-phosphocholine (100µg/mL) (Sigma, DOPC, cat# 850375P), DNA (0.6mg/mL) (Sigma, Herring sperm DNA, cat# D3159), Yeast Synthetic Dropout (4mg/mL) (Sigma, Trp, cat# Y1876), NaCl (51.85mM), MOPS (100mM), KCl (14.94mM), NH_4_Cl (2.28mM), and FeSO_4_ (3.6µM). When preparing ASM, all of the components excluding mucin and FeSO_4_ are dissolved in molecular grade water at a 2x final concentration, pH is adjusted to 6.8. Mucin (Sigma, Mucin from porcine stomach, Type 2) is suspended in water at a 10mg/mL concentration and sterilized by autoclaving. On the day of use, ASM base components are mixed with mucin at a 1 to 1 ratio, subsequently fresh FeSO_4_ stock is prepared and added to the media at a final concentration of 3.6µM.

For the anaerobic growth assays, bacteria were cultured in an anoxic environmental chamber (Coy labs) under atmosphere containing a 5% CO_2_, 5% H_2_, 90% N_2_ gas mixture. After 6h incubation at 37°C, plates were removed from the anoxic chamber, planktonic growth fraction was collected, serially diluted, and plated on Tryptic soy agar plates supplemented with 5% v/v defibrillated sheep blood for enumeration. Biofilm fraction was washed with PBS twice, subsequently 50µL of PBS was added and bacteria was detached from the plastic using a 96-pin replicator. Biofilm fraction was subsequently serially diluted and plated for CFU quantification. Plates for CFU quantification were incubated at 37°C under 5% CO_2_ atmosphere for 18 to 36h until well defined colonies appeared.

### Cell-associated metal content analysis

For the purposes of cell-associated metal content analysis, methodology described previously (69) was adapted. Bacteria from an overnight culture were sub-cultured into tubes containing 10mL TH-YE medium at a staring OD_600_=0.01 and cultured statically for 6h at 37°C at 5% CO_2_. A total of 30 mL of each bacterial culture was collected and pelleted by centrifugation (10min, 4000x*g*, 4°C). Supernatant was discarded and bacteria were subsequently resuspended in Mg- and Ca-free PBS supplemented with 50mM EDTA (pH=7.0). Three washes in EDTA containing PBS were followed by three further washes in PBS. Subsequently, bacterial pellets were frozen and stored at -80°C prior to lipolysis using Labconco FreeZone Benchtop Freeze Dryer. After weighing dry bacterial pellets, these samples were submitted to inductively coupled plasma-mass spectrometry (ICP-MS) analysis at Dartmouth Trace Element Analysis Core. Samples were subjected to nitric acid digestion according to the methodology described previously (70). Concentrations of the following metals were assessed – As, Ba, Ca, Cd, Co, Cu, Fe, K, Mg, Mn, Mo, Ni, Pb, Se, Sr, Zn. Metal content was expressed as ng or µg per mg of dried whole cell pellet.

### Antibiotic susceptibility testing

For antimicrobial sensitivity testing, fresh antibiotic stocks were prepared on the day of testing. Polymyxin B sulfate (Research Products International, cat# 1405-20-5) and colistin sulfate (Sigma, cat# C4461) stocks were prepared directly in TH-YE media. Ciprofloxacin (Sigma, cat# 17850) and levofloxacin (TCI, cat# L0193) stocks were prepared at a 10mg/mL concentration in 0.1N acetic acid. Vancomycin hydrochloride (Sigma, cat# 94747), clindamycin hydrochloride (Research Products International, cat# C41050), and daptomycin (Thermo Scientific, cat# 461371000) stocks were dissolved in molecular grade water. All concentrated antibiotic stocks were sterilized using 0.22µm syringe filter. Antibiotic stocks were added to TH-YE medium to achieve specified final concentrations.

Bacterial strains from overnight cultures were pelleted by centrifugation (6000 x *g*, 3min), and subsequently washed in PBS twice. Next, bacterial OD_600_ was adjusted to 0.02 in TH-YE medium. 96-well plate was filled with 100µL of TH-YE medium containing 2x the desired antibiotic concentration. 100µL of bacteria in TH-YE medium was added to each of the wells resulting in a starting bacterial inoculum of OD_600_=0.01. Medium-only wells were added to allow for background correction in subsequent OD_600_ measurements. Plates were incubated without agitation at 37°C, 5% CO_2_ for 18h. After incubation, bacterial growth was assessed using a Spectra Max M2 plate reader.

### Checkerboard assays

To assess how addition of metal ions impacts bacterial susceptibility to Polymyxin B a checkerboard assay (71) was employed. CaCl_2_, MgCl_2_, ZnSO_4_ stock solutions of 0.5M were prepared in molecular grade water. Polymyxin B solutions were prepared on the day of use by dissolving antibiotic directly in TH-YE medium. Metal and antibiotic stocks were sterilized using a 0.22µm syringe filter. Assays were performed in 96-well plate format. Salt solutions and antibiotics were added to the TH-YE medium and concentrations adjusted by two-fold serial dilutions. Subsequently antibiotic and metal solutions were added to the assay plate in perpendicular dilution series.

Bacterial overnight cultures are pelleted (6000 x *g*, 3min), and washed in PBS two times. Subsequently, bacterial OD_600_ was standardized in TH-YE. Bacteria were inoculated into the checkboard assay plates at an initial OD_600_ equivalent of 0.01 and incubated for 18h at 37°C under 5% CO_2_ atmosphere. After incubation, bacterial growth was assessed using a Spectra Max M2 plate reader. For the biofilm disruption assay, bacteria were grown in TH-YE medium in a 96-well plate, after growth for 18h at 37°C under 5% CO_2_, the medium was removed and replaced with fresh medium containing antibiotic and metals at the specified concentrations. Subsequently, bacteria were cultured for further 6h at 37°C under 5% CO_2_ before assessing bacterial abundance. Following the OD_600_ measurements, bacteria in a plate were disrupted using a 96-pin replicator and plated on tryptic soy agar medium supplemented with 5% sheep blood for a non-quantitative assessment of antibiotic lytic or static inhibitory effect.

Values from OD_600_ measurements, were background corrected against wells containing only medium. Bacterial growth in individual wells was reported relative to untreated control wells, where 1 indicates no change and values approaching 0 correspond to no growth detected.

### Transmission electron microscopy (TEM) imaging

To assess impact of the mutations studied here on *S. sanguinis* cell wall integrity, cells were imaged using transmission electron microscopy (TEM). Bacteria from overnight culture were used to inoculate 10mL TH-YE medium at a starting OD_600_=0.01 and cultured statically for 6h at 37°C, under 5% CO_2_ atmosphere. Subsequently, a total of 40mL of each bacterial culture were pooled into a 50mL centrifuge tube. Bacteria were pelleted by centrifugation at 3000 x *g* for 5min. Supernatant was discarded, and pellet was resuspended in a freshly prepared fixative consisting of glutaraldehyde (2.5%), paraformaldehyde (3.2%), and sodium cacodylate (0.1M, pH7.3). After fixation at room temperature for 1h, bacteria were pelleted, resuspended in fresh fixative and submitted for further fixation, embedding, and imaging at Dartmouth Electron Microscopy Facility. Imaging was done using Thermo Scientific HELIOS 5CX microscope.

For the purposes of cell wall thickness measurements, cells were imaged at 150000x magnification. Cross-sections of >25 individual cells were selected from each sample. Cell wall thickness measurements were performed using ImageJ, each cell was measured at 4-8 locations equally distributed across cell perimeter, avoiding sections in close contact with adjacent cells or sections close to the cell division plane.

### Zeta potential measurements

Procedure for zeta potential measurements was adapted from previous studies (41, 72). Briefly, bacteria were grown in TH-YE medium, sub-cultures were inoculated at an initial OD_600_=0.01 and incubated at 37°C, 5% CO_2_ for 6h. Subsequently, bacteria were pelleted, washed in PBS and normalized to OD_600_=0.1. After bacterial density normalization, cell suspensions were transferred to Malvern Folded Capillary cuvettes. Zeta potential measurements were performed using Zetasizer NanoZS (Malvern Instruments).

### Laurdan membrane fluidity assay

Membrane fluidity was assessed using Laurdan generalized polarization assay. Laurdan is a membrane intercalating fluorescent probe that shifts emission wavelength depending on the amount of water within the membrane, this being indicative of membrane packing and relative fluidity (44, 73, 74). The experimental procedure was adapted from the protocol described previously (46). Bacterial sub-cultures of 10mL TH-YE medium were inoculated at the initial OD_600_ of 0.01 and grown at 37°C, 5% CO_2_ atmosphere for 6h. After incubation bacteria were moved to a 37°C warm room and all subsequent handling and measurement steps were performed at 37°C, and the centrifuge, disposable materials and reagents were pre-warmed before use. 1980µL of bacteria from sub-cultures were transferred to a microcentrifuge tube and 20µL of 1mM Laurdan fluorescent dye (Sigma, cat# 40277) dissolved in dimethylformamide (DMF) was added to each bacterial aliquot. After mixing, bacteria were covered to protect from light and incubated for 5min to allow for dye to integrate into the membrane. Subsequently, bacteria were palleted (7500g, 1min), the supernatant was discarded, and pellet resuspended in Laurdan buffer [137mM NaCl, 2.7mM KCl, 10mM Na_2_HPO_4_, 1.8mM KH_2_PO_4_, 0.2% w/v glucose, 1% v/v DMF, filter sterilized]. Bacteria were washed in Laurdan buffer a total of 4 times. Next, the bacterial density was normalized to an OD_600_=0.4 and three measurement replicates of 100µL were transferred to a 96-well black wall, clear bottom plate. The plate was transferred to a Synergy Neo2 plate reader, fluorescent measurements were performed with excitation at 350nm and emission at 460 and 500nm. OD_600_ measurements were obtained to validate accuracy of the dilution.

To assess treatment impact on membrane fluidity, bacteria were prepared as above, with the exception of OD_600_ adjustment to 0.8. The plate containing bacteria was placed in a plate reader and baseline readings were recorded. Subsequently CaCl_2_, ZnSO_4_, and Polymyxin B treatments were added, the plate was returned to the plate reader for incubation in the dark. Ten min after treatment start, fluorescent readings were recorded to assess treatment impact. Treatment mixtures were prepared as follows: Polymyxin B was dissolved directly into the Laurdan buffer, filter sterilized and diluted to the appropriate concentration. Treatment mixes containing Ca^2+^ or Zn^2+^ ions were prepared by adding the appropriate amount of 0.5M CaCl_2_ or ZnSO_4_ solution to the Laurdan buffer.

Laurdan Generalized Polarization (GP) was calculated using the following formula:

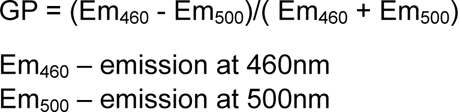

A high GP value is indicative of relatively rigid membrane, while decrease in GP values is associated with increased water content in the membrane, which corresponds to increase in membrane fluidity.

### Fatty acid methyl ester (FAME) analysis

Samples for whole cell fatty methyl ester (FAME) analysis were prepared as follows: Overnight liquid cultures were used to inoculate 150mL TH-YE cultures, which were subsequently incubated statically for 6h at 37°C in 5% CO_2_ atmosphere. Next, bacterial pellets were collected by centrifugation (10min, 4000 x *g*, 4°C) and resuspended in 2mL PBS to allow pellet pooling and transfer to a single microcentrifuge tube. Next bacteria were pelleted by centrifugation (6000 x *g*, 3min, 4°C), the supernatant was discarded, and cell pellets frozen before lipolysis using Labconco FreeZone Benchtop Freeze Dryer. Dried cell pellets were submitted for FAME analysis was performed by Creative Proteomics and subjected to the following extraction protocol.

Samples were weighed into a screw-cap glass vial which contained tritricosanoin as an internal standard (tri-C23:0 TG) (NuCheck Prep, Elysian, MN). A portion of the organic layer was transferred to a screw-cap glass vial and dried in a speed vac. After samples were dried BTM (methanol containing 14% boron trifluoride, toluene, methanol; 35:30:35 v/v/v) (SigmaAldrich, St. Louis, MO) was added. The vial was briefly vortexed and heated in a hot bath at 100°C for 45 minutes. After cooling, hexane (EMD Chemicals, USA) and HPLC grade water was added, the tubes were recapped, vortexed and centrifuged help to separate layers. An aliquot of the hexane layer was transferred to a GC vial. Fatty acids were identified by comparison with a standard mixture of fatty acids (GLC OQ-A, NuCheck Prep, Elysian, MN) which was also used to determine individual fatty acid calibration curves.

## Acknowledgements

This work was supported by National Institutes of Health (R01 AI155424) to G.A.O. The authors also acknowledge the Genomics and Molecular Biology and Trace Element Analysis Shared Resources at the Dartmouth Cancer Center with NCI Cancer Center Support Grant 5P30 CA023108-41. We also thank the Dartmouth Electron Microscopy Facility for their assistance.

## Figure descriptions

Supplementary figure 1 Growth and biofilm formation of the *ecfAAT* mutants in different media conditions.

Planktonic (**A**) and biofilm growth (**B**) of the WT and *ecfAAT* mutants compared in TH-YE and ASM media mixes. In the TH-YE+PBS condition, TH-YE is mixed with PBS at a 1 to 1 ratio. In ASM+TH-YE condition, TH-YE is mixed with ASM at a 1 to 1 ratio. In ASM+PBS condition, ASM is mixed with PBS in a 1 to 1 ratio. CFU counts assessed after 6h static growth under anoxic conditions. Mean and standard deviation of n=4 biological replicates. Statistical analysis using ANOVA with Tukey’s post hoc test, with *, p<0.05, **, p<0.01, ***, p<0.001, ****, p<0.0001.

Supplementary figure 2 ***ecfAAT* mutant complementation eliminates growth defect observed in artificial sputum medium.**

Complementation of the *ecfAAT* mutant strains under planktonic (**A**) and biofilm growth (**B**) compared in TH-YE and ASM media. CFU counts assessed after 6h static growth under anoxic conditions. Mean and standard deviation of n=5 biological replicates. Statistical analysis using ANOVA.

Supplementary figure 3 **The EcfAAT transporter contributes to metal uptake.**

WT and mutant cell metal content assessed by ICP-MS using bacterial grown in TH-YE medium for 6h at 37°C, 5% CO_2_. Cell associated content of arsenic (**A**), barium (**B**), cadmium (**C**), copper (**D**), potassium (**E**), molybdenum (**F**), nickel (**G**), lead (**H**), selenium (**I**), and strontium (**J**). Metal content reported as ng or µg per mg of dry cell weight. Mean and standard deviation of n=3 biological replicates shown. Statistical analysis using ANOVA with Dunnett’s post hoc test, with *, p<0.05, **, p<0.01.

Supplementary figure 4 **Susceptibility of the *ecfAAT* mutants to antibiotic treatment.**

Assessment of WT and *ecfAAT* mutant strain susceptibility to vancomycin (**A**), clindamycin (**B**), ciprofloxacin (**C**), levofloxacin (**D**), and daptomycin (**E**). For this assay, bacteria were inoculated into TH-YE medium containing the specified amount of a given antibiotic. Bacterial growth was assessed through OD_600_ measurements following 18h static growth at 37°C, 5% CO_2_. Mean and standard deviation of n=3 (vancomycin, clindamycin, daptomycin) or n=4 (ciprofloxacin, levofloxacin) biological replicates shown. Statistical analysis using ANOVA.

Supplementary figure 5 **Assessment of the impact of metal supplementation on Polymyxin B efficacy on established biofilms.**

Checkerboard assay assessing how combined metal ion and Polymyxin B exposure impacts a pre-formed WT or Δ*ecfT* mutant biofilm. Both magnesium (0.3125mM to 10mM) (**Panels on left**) and calcium (0.15625mM to 5mM) (**Central panels**) addition protects WT *S. sanguinis* from Polymyxin B toxicity in a dose dependent manner. Supplementation of zinc (0.015625mM to 1mM) (**Panels on right**) increases Polymyxin B toxicity. WT (**Top row**) and Δ*ecfT* (**Bottom row**) mutant biofilm was established in absence of treatment by growing bacteria in TH-YE medium for 18h statically at 37°C, 5% CO_2_, followed by a 6h treatment exposure under these same conditions before optical density measurements were performed. A representative measurement set of a set 3 biological replicates shown, with darker shading indicating higher bacterial amount.

Supplementary figure 6 **A defect in the EcfAAT system does not detectably impact cell wall thickness or overall surface charge.**

Cell wall integrity assessment was performed using transmission electron microscopy (TEM) imaging (**A**). No significant differences in cell wall thickness between WT and mutant strains were seen (**B**). Cell walls of no less than 25 individual cells from each of the strains were measured, with 4 to 8 measurements taken per cell. Statistical analysis using ANOVA. (**C**) Zeta potential measurements were used to assess the overall surface charge for each of the strains. Before analysis, strains were grown in TH-YE medium in 5% CO_2_ atmosphere for 6h. Mean and standard deviation of 3 independent measurements shown. Statistical analysis using ANOVA.

Supplementary figure 7 **Changes in *S. sanguinis* membrane fluidity upon addition polymyxin B with and without supplementation of Ca^2+^ or Zn^2+^ ions.**

Laurdan GP measurements of WT and *ecfAAT* mutants following exposure to Polymyxin B, Ca^2+^ or Zn^2+^ treatments. For the analysis, bacteria were cultured statically, in TH-YE medium, at 37°C, 5% CO_2_. Mean and standard deviation of n=5 biological replicates shown. Statistical analysis using ANOVA with Dunnett’s post hoc test.

Supplementary figure 8 **Changes in EcfAAT mutant membrane composition.**

FAME analysis results showing changes in the relative abundance of individual FAME species, comparing WT and *ecfAAT* mutant strains – (**A**) C16:1n7t; (**B**) C18:1t; (**C**) C18:2n6t; (**D**) C18:2n6; (**E**) C20:0; (**F**) C18:3n6; (**G**) C20:1n9; (**H**) C18:1n9; (**I**) C20:2n6; (**J**) C22:0; (**K**) C20:3n6; (**L**) C20:4n6; (**M**) C24:0; (**N**) C20:5n3; (**O**) C24:1n9; (**P**) C22:4n6; (**Q**) C22:5n6; (**R**) C22:5n3; (**S**) C22:6n3. Individual FAME content also detailed in **Suppl Table 3**. For the purposes of this analysis, bacteria were cultured statically, in TH-YE medium, at 37°C, 5% CO_2_. Mean and standard deviation of n=3 biological replicates shown. Values shown as adjusted to the total FAME content. Statistical analysis using ANOVA with Dunnett’s post hoc test.

## Literature Cited

1. Zhu B, Macleod LC, Kitten T, Xu P. 2018. *Streptococcus sanguinis* biofilm formation & interaction with oral pathogens. Future Microbiol 13:915–932.

2. Martini AM, Moricz BS, Ripperger AK, Tran PM, Sharp ME, Forsythe AN, Kulhankova K, Salgado-Pabón W, Jones BD. 2020. Association of novel *Streptococcus sanguinis* virulence factors with pathogenesis in a native valve infective endocarditis model. Front Microbiol 11:10.

3. Scott JE, O’Toole GA. 2019. The Yin and Yang of streptococcus lung infections in cystic fibrosis: a model for studying polymicrobial interactions. J Bacteriol 201:e00115–19.

4. Kreth J, Merritt J, Shi W, Qi F. 2005. Competition and coexistence between *Streptococcus mutans* and *Streptococcus sanguinis* in the dental biofilm. J Bacteriol 187:7193–7203.

5. Li K, Gifford AH, Hampton TH, O’Toole GA. 2020. Availability of zinc impacts interactions between *Streptococcus sanguinis* and *Pseudomonas aeruginosa* in coculture. J Bacteriol 202:e00618–19.

6. Rogers RR, Kesthely CA, Jean-Pierre F, El Hafi B, O’Toole GA. 2024. Dpr-mediated H2O2 resistance contributes to streptococcus survival in a cystic fibrosis airway model system. Journal of Bacteriology 0:e00176–24.

7. Zhu B, Green SP, Ge X, Puccio T, Nadhem H, Ge H, Bao L, Kitten T, Xu P. 2021. Genome-wide identification of *Streptococcus sanguinis* fitness genes in human serum and discovery of potential selective drug targets. Mol Microbiol 115:658–671.

8. Eitinger T, Rodionov DA, Grote M, Schneider E. 2011. Canonical and ECF-type ATP-binding cassette importers in prokaryotes: diversity in modular organization and cellular functions. FEMS Microbiology Reviews 35:3–67.

9. Rodionov DA, Hebbeln P, Eudes A, ter Beek J, Rodionova IA, Erkens GB, Slotboom DJ, Gelfand MS, Osterman AL, Hanson AD, Eitinger T. 2009. A novel class of modular transporters for vitamins in prokaryotes. Journal of Bacteriology 191:42–51.

10. Rempel S, Stanek WK, Slotboom DJ. 2019. ECF-Type ATP-Binding cassette transporters. Annual Review of Biochemistry 88:551–576.

11. Finkenwirth F, Eitinger T. 2019. ECF-type ABC transporters for uptake of vitamins and transition metal ions into prokaryotic cells. Research in Microbiology 170:358–365.

12. Jochim A, Adolf L, Belikova D, Schilling NA, Setyawati I, Chin D, Meyers S, Verhamme P, Heinrichs DE, Slotboom DJ, Heilbronner S. 2020. An ECF-type transporter scavenges heme to overcome iron-limitation in *Staphylococcus lugdunensis*. eLife 9:e57322.

13. Chatterjee N, Cook LCC, Lyles KV, Nguyen HAT, Devlin DJ, Thomas LS, Eichenbaum Z. 2020. A novel heme transporter from the energy coupling factor family is vital for Group A Dtreptococcus colonization and infections. Journal of Bacteriology 202:10.1128/jb.00205-20.

14. Rodrigo MKD, Saiganesh A, Hayes AJ, Wilson AM, Anstey J, Pickering JL, Iwasaki J, Hillas J, Winslow S, Woodman T, Nitschke P, Lacey JA, Breese KJ, van der Linden MPG, Giffard PM, Tong SYC, Gray N, Stubbs KA, Carapetis JR, Bowen AC, Davies MR, Barnett TC. 2022. Host-dependent resistance of Group A Streptococcus to sulfamethoxazole mediated by a horizontally-acquired reduced folate transporter. Nat Commun 13:6557.

15. Bousis S, Setyawati I, Diamanti E, Slotboom DJ, Hirsch AKH. 2019. Energy-coupling factor transporters as novel antimicrobial targets. Advanced Therapeutics 2:1800066.

16. Diamanti E, Souza PCT, Setyawati I, Bousis S, Monjas L, Swier LJYM, Shams A, Tsarenko A, Stanek WK, Jäger M, Marrink SJ, Slotboom DJ, Hirsch AKH. 2023. Identification of inhibitors targeting the energy-coupling factor (ECF) transporters. Commun Biol 6:1–8.

17. Exapicheidou IA, Shams A, Ibrahim H, Tsarenko A, Backenköhler M, Hamed MM, Diamanti E, Volkamer A, Slotboom DJ, Hirsch AKH. 2024. Hit optimization by dynamic combinatorial chemistry on *Streptococcus pneumoniae* energy-coupling factor transporter ECF-PanT. Chem Commun 60:870–873.

18. Worlitzsch D, Tarran R, Ulrich M, Schwab U, Cekici A, Meyer KC, Birrer P, Bellon G, Berger J, Weiss T, Botzenhart K, Yankaskas JR, Randell S, Boucher RC, Döring G. 2002. Effects of reduced mucus oxygen concentration in airway *Pseudomonas* infections of cystic fibrosis patients. J Clin Invest 109:317–325.

19. Schobert M, Jahn D. 2010. Anaerobic physiology of *Pseudomonas aeruginosa* in the cystic fibrosis lung. International Journal of Medical Microbiology 300:549–556.

20. Ledger EVK, Sabnis A, Edwards AM. 2022. Polymyxin and lipopeptide antibiotics: membrane-targeting drugs of last resort. Microbiology (Reading) 168:001136.

21. Ayoub Moubareck C. 2020. Polymyxins and bacterial membranes: a review of antibacterial activity and mechanisms of resistance. Membranes (Basel) 10:181.

22. Morrison DC, Jacobs DM. 1976. Binding of Polymyxin B to the lipid A portion of bacterial lipopolysaccharides. Immunochemistry 13:813–818.

23. Manioglu S, Modaresi SM, Ritzmann N, Thoma J, Overall SA, Harms A, Upert G, Luther A, Barnes AB, Obrecht D, Müller DJ, Hiller S. 2022. Antibiotic polymyxin arranges lipopolysaccharide into crystalline structures to solidify the bacterial membrane. Nat Commun 13:6195.

24. Port GC, Vega LA, Nylander AB, Caparon MG. 2014. *Streptococcus pyogenes* Polymyxin B-Resistant Mutants Display Enhanced ExPortal Integrity. J Bacteriol 196:2563–2577.

25. Rudilla H, Pérez-Guillén I, Rabanal F, Sierra JM, Vinuesa T, Viñas M. 2018. Novel synthetic polymyxins kill Gram-positive bacteria. Journal of Antimicrobial Chemotherapy 73:3385–3390.

26. Khadka NK, Aryal CM, Pan J. 2018. Lipopolysaccharide-dependent membrane permeation and lipid clustering caused by cyclic lipopeptide colistin. ACS Omega 3:17828–17834.

27. Moore RA, Bates NC, Hancock RE. 1986. Interaction of polycationic antibiotics with *Pseudomonas aeruginosa* lipopolysaccharide and lipid A studied by using dansyl-polymyxin. Antimicrob Agents Chemother 29:496–500.

28. Nikaido H. 2003. Molecular basis of bacterial outer membrane permeability revisited. Microbiol Mol Biol Rev 67:593–656.

29. D’Amato RF, Thornsberry C, Baker CN, Kirven LA. 1975. Effect of calcium and magnesium Ions on the susceptibility of *Pseudomonas* species to tetracycline, gentamicin, Polymyxin B, and Carbenicillin. Antimicrob Agents Chemother 7:596–600.

30. Yu Z, Qin W, Lin J, Fang S, Qiu J. 2015. Antibacterial mechanisms of polymyxin and bacterial resistance. BioMed Research International 2015:e679109.

31. De Oliveira DMP, Bohlmann L, Conroy T, Jen FE-C, Everest-Dass A, Hansford KA, Bolisetti R, El-Deeb IM, Forde BM, Phan M-D, Lacey JA, Tan A, Rivera-Hernandez T, Brouwer S, Keller N, Kidd TJ, Cork AJ, Bauer MJ, Cook GM, Davies MR, Beatson SA, Paterson DL, McEwan AG, Li J, Schembri MA, Blaskovich MAT, Jennings MP, McDevitt CA, von Itzstein M, Walker MJ. 2020. Repurposing a neurodegenerative disease drug to treat Gram-negative antibiotic-resistant bacterial sepsis. Science Translational Medicine 12:eabb3791.

32. Rihacek M, Kosaristanova L, Fialova T, Kuthanova M, Eichmeier A, Hakalova E, Cerny M, Berka M, Palkovicova J, Dolejska M, Svec P, Adam V, Zurek L, Cihalova K. 2023. Zinc effects on bacteria: insights from *Escherichia coli* by multi-omics approach. mSystems 8:e00733–23.

33. Xia P, Lian S, Wu Y, Yan L, Quan G, Zhu G. 2021. Zinc is an important inter-kingdom signal between the host and microbe. Vet Res 52:39.

34. LaPorte DC, Rosenthal KS, Storm DR. 1977. Inhibition of *Escherichia coli* growth and respiration by polymyxin B covalently attached to agarose beads. Biochemistry 16:1642–1648.

35. Xiong YQ, Mukhopadhyay K, Yeaman MR, Adler-Moore J, Bayer AS. 2005. Functional interrelationships between cell membrane and cell wall in antimicrobial peptide-mediated killing of *Staphylococcus aureus*. Antimicrobial Agents and Chemotherapy 49:3114–3121.

36. Percy MG, Gründling A. 2014. Lipoteichoic acid synthesis and function in Gram-positive bacteria. Annual Review of Microbiology 68:81–100.

37. Brown S, Santa Maria JP, Walker S. 2013. Wall teichoic acids of Gram-positive bacteria. Annu Rev Microbiol 67:10.1146/annurev-micro-092412-155620.

38. Kamar R, Réjasse A, Jéhanno I, Attieh Z, Courtin P, Chapot-Chartier M-P, Nielsen-Leroux C, Lereclus D, el Chamy L, Kallassy M, Sanchis-Borja V. 2017. DltX of *Bacillus thuringiensis* Is essential for d-alanylation of teichoic acids and resistance to antimicrobial response in insects. Front Microbiol 8:1437.

39. Yin J, Meng Q, Cheng D, Fu J, Luo Q, Liu Y, Yu Z. 2020. Mechanisms of bactericidal action and resistance of polymyxins for Gram-positive bacteria. Appl Microbiol Biotechnol 104:3771–3780.

40. Sohlenkamp C, Galindo-Lagunas KA, Guan Z, Vinuesa P, Robinson S, Thomas-Oates J, Raetz CRH, Geiger O. 2007. The lipid lysyl-phosphatidylglycerol is present in membranes of *Rhizobium tropici* CIAT899 and confers increased resistance to Polymyxin B under acidic growth conditions. MPMI 20:1421–1430.

41. Ayala-Torres C, Hernández N, Galeano A, Novoa-Aponte L, Soto C-Y. 2014. Zeta potential as a measure of the surface charge of mycobacterial cells. 3. Ann Microbiol 64:1189–1195.

42. Wilson WW, Wade MM, Holman SC, Champlin FR. 2001. Status of methods for assessing bacterial cell surface charge properties based on zeta potential measurements. Journal of Microbiological Methods 43:153–164.

43. Mohapatra SS, Dwibedy SK, Padhy I. 2021. Polymyxins, the last-resort antibiotics: Mode of action, resistance emergence, and potential solutions. J Biosci 46:85.

44. Orlikowska-Rzeznik H, Krok E, Chattopadhyay M, Lester A, Piatkowski L. 2023. Laurdan discerns lipid membrane hydration and cholesterol content. J Phys Chem B 127:3382–3391.

45. Sanchez SA, Tricerri MA, Gratton E. 2012. Laurdan generalized polarization fluctuations measures membrane packing micro-heterogeneity in vivo. Proceedings of the National Academy of Sciences 109:7314–7319.

46. Wenzel M, Vischer NOE, Strahl H, Hamoen LW. 2018. Assessing membrane fluidity and visualizing fluid membrane domains in bacteria using fluorescent membrane dyes. Bio Protoc 8:e3063.

47. Santos DES, Pol-Fachin L, Lins RD, Soares TA. 2017. Polymyxin binding to the bacterial outer membrane reveals cation displacement and increasing membrane curvature in susceptible but not in resistant lipopolysaccharide chemotypes. J Chem Inf Model 57:2181–2193.

48. Ginez LD, Osorio A, Vázquez-Ramírez R, Arenas T, Mendoza L, Camarena L, Poggio S. 2022. Changes in fluidity of the *E. coli* outer membrane in response to temperature, divalent cations and Polymyxin B show two different mechanisms of membrane fluidity adaptation. The FEBS Journal 289:3550–3567.

49. Zavascki AP, Goldani LZ, Li J, Nation RL. 2007. Polymyxin B for the treatment of multidrug-resistant pathogens: a critical review. Journal of Antimicrobial Chemotherapy 60:1206–1215.

50. Pristovšek P, Kidrič J. 1999. Solution structure of Polymyxins B and E and effect of binding to lipopolysaccharide: An NMR and Molecular Modeling Study. J Med Chem 42:4604–4613.

51. Shatri G, Tadi P. 2024. Polymyxin StatPearls. StatPearls Publishing, Treasure Island (FL).

52. Trimble MJ, Mlynárčik P, Kolář M, Hancock REW. 2016. Polymyxin: Alternative mechanisms of action and resistance. Cold Spring Harb Perspect Med 6:a025288.

53. Nicas TI, Hancock REW. 1983. Alteration of susceptibility to EDTA, Polymyxin B and gentamicin In *Pseudomonas aeruginosa* by divalent cation regulation of outer membrane protein H1. Microbiology 129:509–517.

54. Davis SD, Iannetta A, Wedgwood RJ. 1971. Activity of colistin *against Pseudomonas aeruginosa*: Inhibition by calcium. The Journal of Infectious Diseases 124:610–612.

55. Landman D, Georgescu C, Martin DA, Quale J. 2008. Polymyxins Revisited. Clin Microbiol Rev 21:449–465.

56. Thomas KJ, Rice CV. 2014. Revised model of calcium and magnesium binding to the bacterial cell wall. Biometals 27:1361–1370.

57. Velkov T, Thompson PE, Nation RL, Li J. 2010. Structure—activity relationships of polymyxin antibiotics. J Med Chem 53:1898–1916.

58. Schleimer N, Kaspar U, Drescher M, Seggewiß J, von Eiff C, Proctor RA, Peters G, Kriegeskorte A, Becker K. 2018. The energy-coupling factor transporter module EcfAA’T, a novel candidate for the genetic basis of fatty acid-auxotrophic small-colony variants of *Staphylococcus aureus*. Front Microbiol 9.

59. Xu P, Ge X, Chen L, Wang X, Dou Y, Xu JZ, Patel JR, Stone V, Trinh M, Evans K, Kitten T, Bonchev D, Buck GA. 2011. Genome-wide essential gene identification in *Streptococcus sanguinis*. Sci Rep 1:125.

60. Luke R. Joyce, Ziqiang Guan, Kelli L. Palmer. 2019. Phosphatidylcholine biosynthesis in Mitis group streptococci via host metabolite scavenging. Journal of Bacteriology 201.

61. Freeman CD, Hansen T, Urbauer R, Wilkinson BJ, Singh VK, Hines KM. 2024. Defective *pgsA* contributes to increased membrane fluidity and cell wall thickening in *Staphylococcus aureus* with high-level daptomycin resistance. mSphere 9:e00115–24.

62. Kiefer AF, Bousis S, Hamed MM, Diamanti E, Haupenthal J, Hirsch AKH. 2022. Structure-guided optimization of small-molecule folate uptake inhibitors targeting the energy-coupling factor transporters. J Med Chem 65:8869–8880.

63. Bousis S, Winkler S, Haupenthal J, Fulco F, Diamanti E, Hirsch AKH. 2022. An efficient way to screen inhibitors of Energy-Coupling Factor (ECF) transporters in a bacterial uptake assay. Int J Mol Sci 23:2637.

64. Drost M, Diamanti E, Fuhrmann K, Goes A, Shams A, Haupenthal J, Koch M, Hirsch AKH, Fuhrmann G. 2022. Bacteriomimetic liposomes improve antibiotic activity of a novel energy-coupling factor transporter inhibitor. Pharmaceutics 14.

65. Rhodes DV, Crump KE, Makhlynets O, Snyder M, Ge X, Xu P, Stubbe J, Kitten T. 2014. Genetic characterization and role in virulence of the ribonucleotide reductases of *Streptococcus sanguinis*. J Biol Chem 289:6273–6287.

66. Paik S, Senty L, Das S, Noe JC, Munro CL, Kitten T. 2005. Identification of virulence determinants for endocarditis in *Streptococcus sanguinis* by signature-tagged mutagenesis. Infect Immun 73:6064–6074.

67. Turner KH, Wessel AK, Palmer GC, Murray JL, Whiteley M. 2015. Essential genome of *Pseudomonas aeruginosa* in cystic fibrosis sputum. Proc Natl Acad Sci U S A 112:4110–4115.

68. Clay ME, Hammond JH, Zhong F, Chen X, Kowalski CH, Lee AJ, Porter MS, Hampton TH, Greene CS, Pletneva EV, Hogan DA. 2020. *Pseudomonas aeruginosa lasR* mutant fitness in microoxia is supported by an Anr-regulated oxygen-binding hemerythrin. Proc Natl Acad Sci U S A 117:3167–3173.

69. Turner AG, Ong CY, Gillen CM, Davies MR, West NP, McEwan AG, Walker MJ. 2015. Manganese homeostasis in Group A Streptococcus is critical for resistance to oxidative stress and virulence. mBio 6:10.1128/mbio.00278-15.

70. Heck JE, Andrew AS, Onega T, Rigas JR, Jackson BP, Karagas MR, Duell EJ. 2009. Lung cancer in a U.S. population with low to moderate arsenic exposure. Environmental Health Perspectives 117:1718–1723.

71. Bellio P, Fagnani L, Nazzicone L, Celenza G. 2021. New and simplified method for drug combination studies by checkerboard assay. Methods X 8.

72. Kłodzińska E, Szumski M, Dziubakiewicz E, Hrynkiewicz K, Skwarek E, Janusz W, Buszewski B. 2010. Effect of zeta potential value on bacterial behavior during electrophoretic separation. Electrophoresis 31:1590–1596.

73. Sánchez SA, Tricerri MA, Gunther G, Gratton E. 2007. Laurdan generalized polarization: from cuvette to microscope. Modern research and educational topics in microscopy 2:1007–1014.

74. Parasassi T, Gratton E. 1995. Membrane lipid domains and dynamics as detected by Laurdan fluorescence. J Fluoresc 5:59–69.

